# Vina-GPU 2.1: towards further optimizing docking speed and precision of AutoDock Vina and its derivatives

**DOI:** 10.1101/2023.11.04.565429

**Authors:** Shidi Tang, Ji Ding, Xiangyu Zhu, Zheng Wang, Haitao Zhao, Jiansheng Wu

## Abstract

AutoDock Vina and its derivatives have established themselves as a prevailing pipeline for virtual screening in contemporary drug discovery. Our Vina-GPU method leverages the parallel computing power of GPUs to accelerate AutoDock Vina, and Vina-GPU 2.0 further enhances the speed of AutoDock Vina and its derivatives. Given the prevalence of large virtual screens in modern drug discovery, the improvement of speed and accuracy in virtual screening has become a longstanding challenge. In this study, we propose Vina-GPU 2.1, aimed at enhancing the docking speed and precision of AutoDock Vina and its derivatives through the integration of novel algorithms to facilitate improved docking and virtual screening outcomes. Building upon the foundations laid by Vina-GPU 2.0, we introduce a novel algorithm, namely Reduced Iteration and Low Complexity BFGS (RILC-BFGS), designed to expedite the most time-consuming operation. Additionally, we implement grid cache optimization to further enhance the docking speed. Furthermore, we employ optimal strategies to individually optimize the structures of ligands, receptors, and binding pockets, thereby enhancing the docking precision. To assess the performance of Vina-GPU 2.1, we conduct extensive virtual screening experiments on three prominent targets, utilizing two fundamental compound libraries and seven docking tools. Our results demonstrate that Vina-GPU 2.1 achieves an average 4.97-fold acceleration in docking speed and an average 342% improvement in EF1% compared to Vina-GPU 2.0. The source code and tools for Vina-GPU 2.1 are freely available at https://github.com/DeltaGroupNJUPT/Vina-GPU-2.1, accompanied by comprehensive instructions and illustrative examples.

## I. Introduction

Molecular docking plays a pivotal role in elucidating the binding interactions between different molecular entities, such as a drug and its target. As a result, it has gained significant popularity in modern drug discovery endeavors [1]. This computational technique facilitates the prediction of proteinligand interactions, enabling the identification of energetically favorable conformations in the bound state. In addition, molecular docking tools provide an efficient and cost-effective means for the initial stages of drug design, allowing the identification of potential compounds and the evaluation of their binding affinities [2], [3], [4]. Among the available tools, the AutoDock Vina suite and its derivatives are widely regarded as the preferred choice for molecular docking, owing to their remarkable speed and accuracy [5].

Previous investigations have revealed that early iterations of AutoDock Vina and its derivatives are incapable of meeting the speed requirements imposed by contemporary drug discovery practices, necessitating the development of accelerated versions [5], [6]. Graphics Processing Units (GPUs) have emerged as an ideal solution for common users due to their accessibility, cost-effectiveness, and ease of implementation. Leveraging the power of GPUs, our Vina-GPU method addresses the challenge of parallelizing the AutoDock Vina suite on GPUs, which is hindered by the serial design of the AutoDock Vina algorithm. As a result, we achieve a notable 14-fold acceleration in a typical virtual screening scenario [6]. Moreover, in a benchmark dataset comprising 140 complexes, Vina-GPU demonstrates an average docking acceleration of 21-fold and a maximum acceleration of 50-fold, while maintaining docking accuracy comparable to the original AutoDock Vina[7].

To further enhance the speed of the AutoDock Vina suite, we introduce the Vina-GPU 2.0 method, which encompasses new docking algorithms (QuickVina 2 and QuickVina-W) designed for GPUs [5]. Due to the disparities in their respective docking algorithms, Vina-GPU 2.0 employs distinct strategies for GPU acceleration [5]. When conducting virtual screening experiments on two protein kinase targets, namely RIPK1 and RIPK3, sourced from the DrugBank database, our Vina-GPU 2.0 method achieves remarkable average docking accelerations of 65.6-fold, 1.4-fold, and 3.6-fold when compared to the original AutoDock Vina, QuickVina 2, and QuickVina-W, respectively, while ensuring comparable docking accuracy [5]. Regarding the acceleration of virtual screening, there remains scope for further improvement in Vina-GPU 2.0. Notably, the implementation of the Broyden–Fletcher–Goldfarb–Shanno (BFGS) optimization algorithm constitutes the most time-consuming operation (as shown in Table II) during the ligand conformation update process in docking. Thus, it is imperative to devise a more efficient BFGS optimization algorithm that is well-suited for the demands of virtual screening under realworld conditions. Additionally, optimizing the sharing of grid cache can yield further acceleration benefits for QuickVina 2-GPU and QuickVina-W-GPU, particularly by eliminating the need to recalculate the grid cache for each ligand in a virtual ligand screening scenario [5]. In virtual screening, the identification of promising compounds stands as the primary objective, and enhancing screening accuracy or minimizing the loss of best-hit compounds is of paramount importance [8]. Numerous approaches have been proposed to improve screening accuracy, focusing primarily on expanding the conformational search space [9] and optimizing the scoring function [10] during molecular docking. Moreover, our virtual screening experiments have highlighted the potential for enhancing screening accuracy through the improvement of the threedimensional structure quality of ligand and receptor molecules, as well as the data quality of docked pocket structures [11], [12]. These aspects warrant careful attention as they hold promise for augmenting screening accuracy.

Continuing from Vina-GPU 2.0, the development of Vina-GPU 2.1 focuses on further enhancing docking speed and accuracy to improve the overall performance of docking and virtual screening. **To optimize docking speed**, we introduce a new algorithm called Reduced Iteration and Low Complexity BFGS (RILC-BFGS), specifically designed to accelerate the most time-consuming operation, BFGS (as shown in Table II). RILC-BFGS reduces data storage space, communication, and computation costs by only storing the latest iterations, resulting in significant speed improvements [13]. Additionally, we implement Grid Cache optimization in QuickVina 2-GPU [5] and QuickVina-W-GPU [5]. This optimization minimizes communication costs between the CPU and GPU by eliminating the need to recalculate the grid cache for each ligand, thereby speeding up the molecular docking process [5]. **To enhance docking accuracy**, we employ different strategies to optimize the structures of receptors, binding pockets, and ligands. For receptors, we utilize a cross-docking scheme to identify the most suitable cocrystal structure for docking during the screening process [11]. This step helps ensure that the appropriate receptor structure is used for accurate docking. For binding pockets, we employ the COACH-D server, which employs molecular docking to refine ligand binding poses. This refinement process improves the prediction of protein-ligand binding sites, leading to enhanced docking accuracy [14]. Finally, for ligands, we utilize the open-source program Gypsum-DL to prepare small-molecule libraries with highquality 3D coordinates. This preparation includes considering feasible ionization, tautomeric, and ring-conformational variants, resulting in improved ligand structures for docking [12]. These combined strategies contribute to the overall enhancement of docking accuracy in Vina-GPU 2.1.

In the pursuit of better docking performance, benchmarking studies on a set of 140 complexes demonstrate the effectiveness of Vina-GPU 2.1. It achieves an average docking speed acceleration of 1.62-fold, 6.89-fold, and 3.56-fold compared to the original Vina-GPU+, QuickVina2-GPU, and QuickVinaW-GPU methods from Vina-GPU 2.0, while maintaining comparable docking accuracy [5]. This showcases the improved efficiency and speed of Vina-GPU 2.1 in the docking process. **For enhanced virtual screening**, extensive experiments were conducted on three high-profile targets using two fundamental compound libraries and seven different docking tools with various experimental configurations. The results reveal that Vina-GPU 2.1 achieves an average screening speed acceleration of 2.18-fold, 8.34-fold, and 4.42-fold when compared to Vina-GPU+, QuickVina2-GPU, and QuickVina-W-GPU methods, respectively, when screening on the DrugBank library and Selleck library. Additionally, Vina-GPU 2.1 demonstrates average improvements of 342% improvement in EF1% (Enrichment Factor at 1%) against Vina-GPU 2.0. Among the components of Vina-GPU 2.1, QuickVina2-GPU 2.1 exhibits the fastest average screening speed, while AutoDockVina-GPU 2.1 showcases the best average screening accuracy, with a 57% TOP1 RMSD (Root Mean Square Deviation) of less than 2 Å [5]. Ablation experiments further confirm that the optimization of receptor, pocket, and ligand structures all contribute to enhancing docking accuracy in most cases. Notably, the novel Reduced Iteration and Low Complexity BFGS (RILC-BFGS) algorithm achieves an average improvement of 47.27% in screening speed compared to the original BFGS algorithm, while maintaining comparable screening accuracy [13]. To facilitate the usage of Vina-GPU 2.1, the codes and tools, along with explicit instructions and examples, are available at https://github.com/DeltaGroupNJUPT/Vina-GPU-2.1. This allows researchers and users to access and utilize Vina-GPU 2.1 effectively.

## II. Methodology

### A. Overall architecture

Figure 1 illustrates the overall architecture of Vina-GPU 2.1, which involves four steps of optimization in both the screening pipeline and docking algorithm. The first step focuses on selecting the best structure of the target receptor using the cross-docking method [11]. If necessary, the second step involves identifying the potential binding pocket of the selected structure using the COACH-D method [14]. In the third step, molecules in the databases are converted from 2D to 3D using the efficient Gypsum-DL method [12]. Finally, the fourth step entails the novel RILC-BFGS algorithm, which consists of four parts for the molecular docking process. The first three steps contribute to improved accuracy in virtual screening, as described in Section II D. The last step aims to further accelerate the docking process using the novel RILC-BFGS and GCS methods, as explained in II B and II C. The RILC-BFGS method optimizes the time-consuming BFGS operation (outlined in Table II), while the GCS method optimizes redundancies in the original docking procedure.

**Fig. 1.**
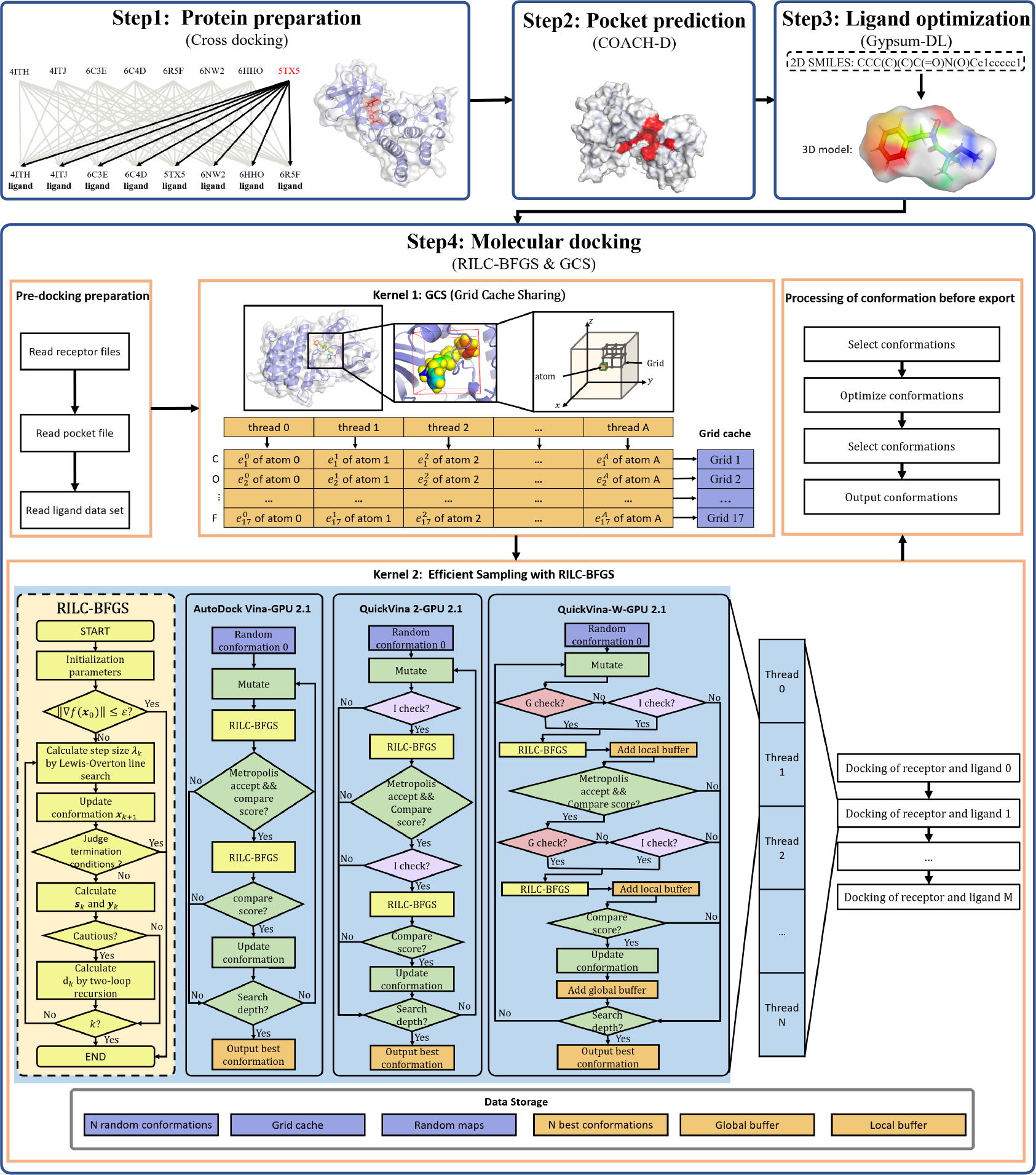
The Vina-GPU 2.1 architecture. It comprises four steps including protein preparation, pocket preparation, ligand preparation and molecular docking with RILC-BFGS and GCS methods.

In the fourth step of Molecular docking, it can be further divided into three parts. The first part involves pre-processing, where the receptor, pocket, and ligand dataset are read and prepared for the docking process. The second part, referred to as “kernel 1” computation, calculates the grid caches of the binding pocket. This computation is performed in parallel using multi-threading with the OpenCL kernel function. The third part, referred to as “kernel 2” computation, corresponds to the actual molecular docking. It involves using three alternative methods: AutoDock Vina-GPU 2.1, QuickVina 2-GPU 2.1, and QuickVina-W-GPU 2.1. All three methods employ the novel RILC-BFGS algorithm for the docking process.

In the fourth step of molecular docking, the AutoDock Vina-GPU 2.1 method follows a specific process. First, it generates *N* initial ligand conformations, with each conformation corresponding to an OpenCL thread. These conformations undergo random mutations in terms of position, orientations, and torsions. Then, the novel RILC-BFGS algorithm is applied to optimize the conformations, and binding scores are calculated for each conformation. A Metropolis acceptance criterion is used to determine whether to accept the optimized conformation. If the criterion is met, another RILC-BFGS iteration is performed, updating the ligand conformation. If not, the next iteration is initiated. After a certain number of iterations, each OpenCL thread outputs the best conformation obtained during the iterations. The QuickVina 2-GPU 2.1 and QuickVina-WGPU 2.1 methods share a similar process with AutoDock Vina-GPU 2.1. However, they include additional steps: I-check (for QuickVina 2-GPU 2.1 and QuickVina-W-GPU 2.1) and G-check (for QuickVina-W-GPU 2.1). These steps enable the skipping of unnecessary iterations. Notably, the G-check process requires additional local and global buffers to store intermediate variables. In the final part, the conformations from all *N* threads are gathered, and the best conformations are selected for further optimization. The final ligand files are then exported. This detailed description outlines the specific steps and variations within the molecular docking process for each method.

### B. Efficient sampling with RILC-BFGS

The sampling method of AutoDock Vina and its derivatives (QuickVina 2 and QuickVina-W) is described as the iterated local search global optimizer (ILSGO) method [7], which comprises a simulating annealing method (global optimizer) and an embedded Broyden–Fletcher

–Goldfarb–Shanno (BFGS) algorithm (local search). Our Vina-GPU [6] and Vina-GPU 2.0 [5] proposed a parallel version of ILSGO method and implemented it into the graph processing unit (GPU). Although ILSGO has shown high accuracy and efficiency in the last decades, more optimizations are still needed in both accuracy and speed for the large virtual screening scenario nowadays. Basically, the BFGS algorithm in ILSGO includes the calculation of the docking energy *f* of the binding conformation *C*, the respective derivatives *g* of the scoring function and an iteration of updating the approximate matrix *H*. Within the iteration, an Armijo-Goldstein approach is adopted to conduct the line search process, which finds the best conformation along with the direction *d* that the scoring function descends.

According to our observation, BFGS local search method is the performance bottleneck of ILSGO and inefficient. This deficiency is caused by the following three aspects: (1) **long iteration**. The outer loop in the BFGS method and the inner loop in the Armijo-Goldstein approach are extremely time-consuming; (2) **inaccurate line search**. The adopted Armijo-Goldstein line search is a simple method with many limitations like low convergence and accuracy[15]; (3) **high computational complexity**. The matrix update step in the BFGS method is computationally intensive with a quadratic complexity (*O*(*n*^2^)), which hampers the speed of optimization. To alleviate these aspects, we propose a novel **r**educed **i**teration and **l**ow **c**omplexity BFGS algorithm named RILCBFGS (Algorithm 1). The RILC-BFGS method consists of three modifications aimed at the above three aspects accordingly. **First**, we adopted an early stop criterion (Line 18-20 Algorithm 1) given by *g*^*T*^ *d <* 0 to reduce the long iterations. If the criterion is not satisfied, which indicates that the direction *d* is not pointing to the direction that descending the scoring function, the calculations afterwards are considered not meaningful and the RILC-BFGS process will be terminated, thus, accelerate the iteration process. Besides, in each line search process, we abandon the conformation whose energy is higher than the original (before line search) conformations’ when the max iteration is reached. **Second**, we replace the Armijo-Goldstein line search with an inexact Lewis-Overton line search method[16], which is more robust for local minimum and more accurate for non-convex objective function like the AutoDock Vina scoring function. The Lewis-Overton line search method (Algorithm 2) considers both Armijo-Goldstein (Line 7 of Algorithm 2) and weak Wolfe condition [17] (Line 9 of Algorithm 2) and calculate the step *λ* through a maximum of *L* iterations. **Third**, we adopted a low complexity LBFGS[18] algorithm with two-loop recursion and cautious updating[15] to replace the previous BFGS algorithm. The two-loop recursion is given in Algorithm 3, in which two memory matrices *S* and *Y* are used to store the historical *s*_*i*_ (Equation. 1) and *y*_*i*_ (Equation. 2) (*i* = 1, 2, …, *m*) during iterations.*m* represents the capacity of the memory matrix. The cautious updating is given by Equation. 3, which determines whether to update the descending direction *d* on the consideration of *s*_*i*_ and *y*_*i*_.

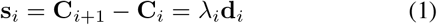

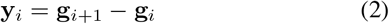

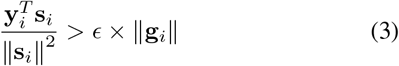

The overall algorithm of RILC-BFGS is given in Algorithm 1. The inputs are conformation **C**_0_, the memory matrix capacity *m*, the maximum iteration count *K* and the gradient of the scoring function under **C**_0_. In each of *K* iterations, RILC-BFGS first calculates the gradient *g* and then obtains the updating step *λ* and the updated conformation **C** (Line 3-6 of Algorithm 1) with the Lewis-Overton line search (Algorithm 2). Next, we compare the energy that before and after the line search, if the energy does not decline, we reset the conformation to the previous (before the line search) one (Line 7-9 of Algorithm 1). If the gradient **g** is small enough (i.e., less than 1*e*^*-*5^), then we terminate the iteration (Line 11-13 of Algorithm 1), otherwise we calculate *s* and *y* based on Equation. 1 and Equation. 2 (Line 14 of Algorithm 1). Finally, we update *d* if cautious condition is satisfied (Line 15-17 of Algorithm 1). Notably, the memory matrices *S* and *Y* belong to *R*^*m×n*^. As a result, for scenarios where *m* is relatively small, the computational complexity of the RILC-BFGS algorithm is *O*(*mn*), which is lower than that of the BFGS algorithm (*O*(*n*^2^)), rendering it computationally more efficient.

### C. Compact workflow with GCS (grid cache sharing)

Vina-GPU 2.1 incorporates a **G**rid **C**ache **S**haring (GCS) mechanism, as depicted in Figure 1 (step 4, Kernel 1). This mechanism was also implemented in Vina-GPU+ [5]. GCS facilitates the shared utilization of the grid cache for each ligand calculation during virtual screening. By avoiding redundant calculations of the grid cache for each ligand, GCS significantly reduces the overall docking runtime. The GCS method is a general technique that can be employed in any grid-cache-like docking algorithm. In the case of Vina-GPU 2.1, QuickVina2-GPU 2.1, and QuickVina-W-GPU 2.1, we have also applied the GCS method. By implementing GCS, these algorithms can take advantage of the shared use of the grid cache, resulting in improved efficiency and reduced computational time during the docking process (as described in III E).

### D. Structure optimization

In the previous Vina-GPU 2.0, we conducted typical methods to prepare the structures of the target receptor, binding pocket and the molecules. However, these typical methods are not often the best in the scenario of virtual screening. Therefore, in Vina-GPU 2.1, we try to optimize these structures in the following three steps (receptor preparation, binding pocket prediction and ligand optimization) and help to improve the accuracy of the virtual screening with Vina-GPU 2.1.

#### Algorithm 1 RILC-BFGS

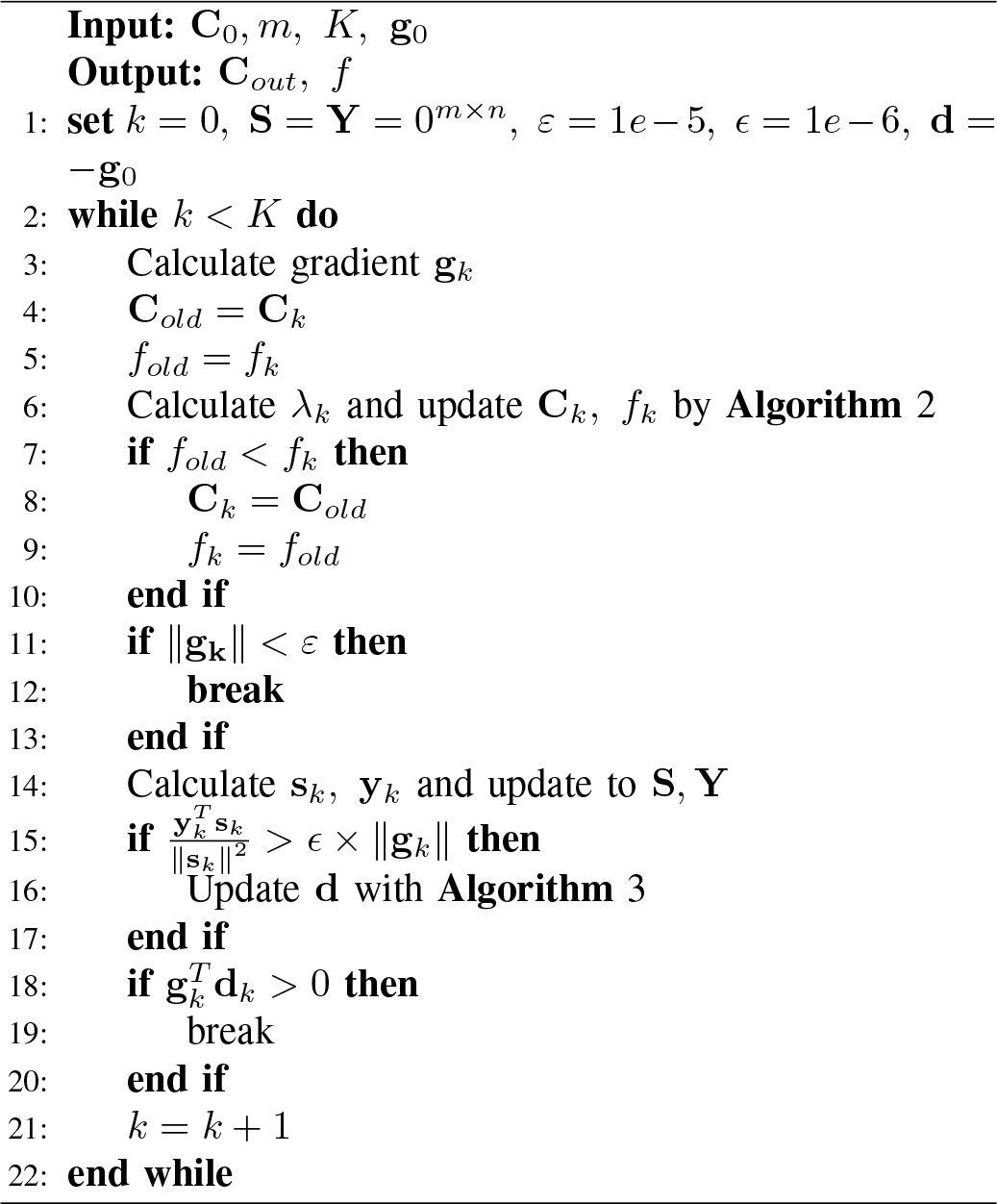

#### Algorithm 2 Lewis-Overton line search of RILC-BFGS

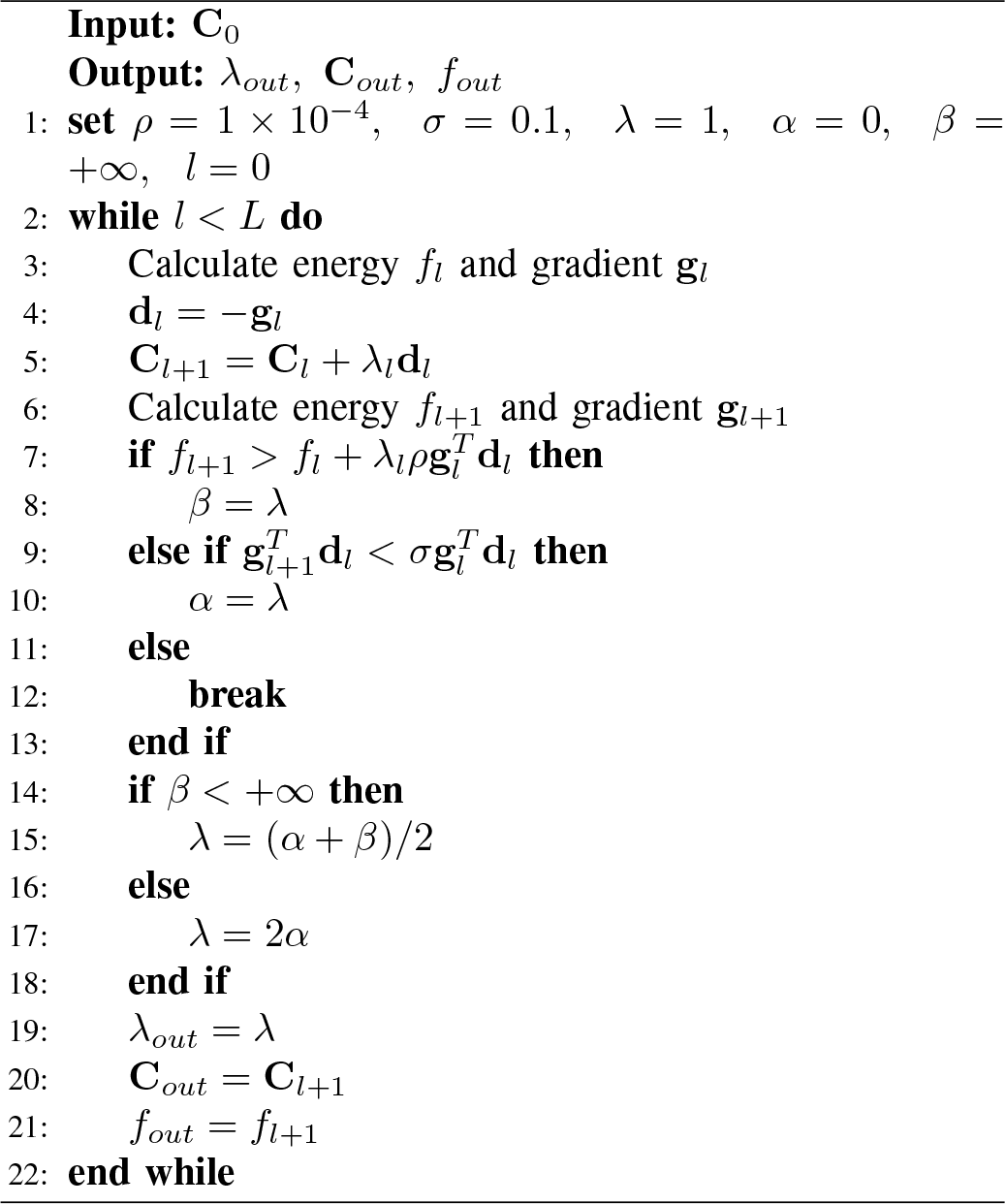

#### Algorithm 3 two-loop recursion of RILC-BFGS

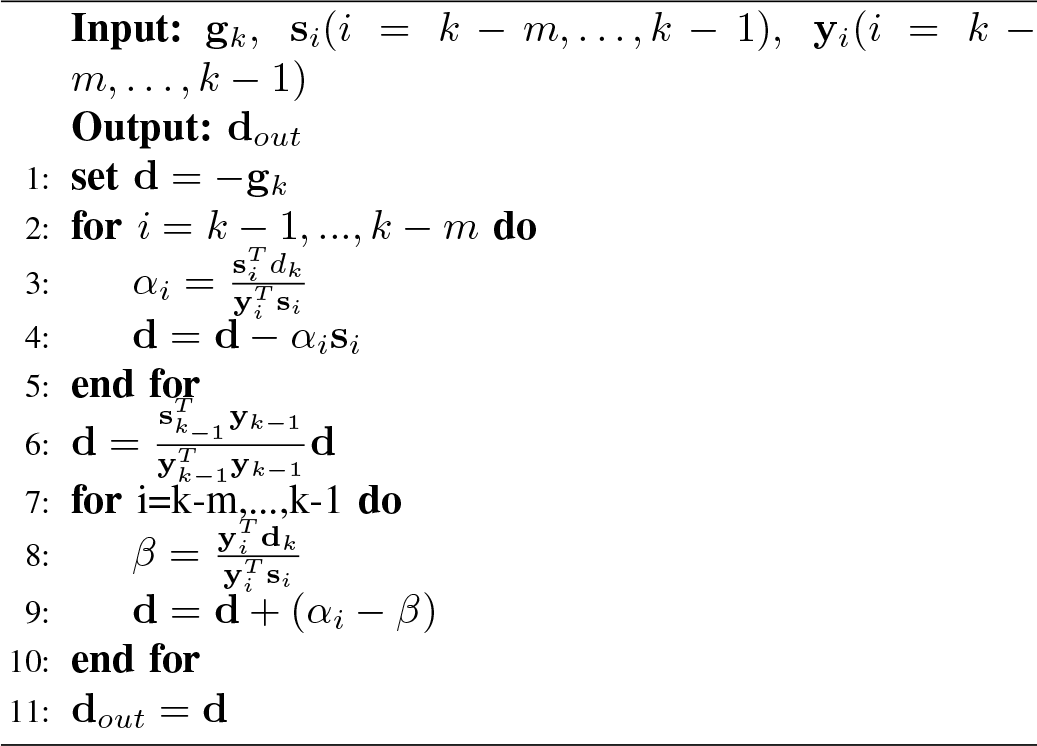

#### 1) Receptor Preparation

The selection and preparation of the receptor is the first and crucial step in structure-based virtual screening, as the structural quality, particularly the resolution of the crystal structure, significantly impacts the accuracy of the screening process. In the conventional approach utilized in Vina-GPU 2.0, all available receptor structures (PDB IDs) corresponding to the target under investigation are obtained from the Protein Data Bank [19]. Typically, the structure with the lowest resolution is chosen, as it is commonly believed that lower resolution corresponds to increased structural accuracy. However, it has been observed that the receptor structure selected using this method does not consistently yield optimal screening accuracy. To address this limitation, Vina-GPU 2.1 incorporates an alternative cross-docking method [11], which offers improved selection through cross-validation. In this approach, each receptor structure is individually docked with ligands extracted from other structures (different PDB IDs). Subsequently, the docking results are assessed using the RootMean-Square-Distance (RMSD) criterion, which measures the deviation between the ligand conformations obtained from the original structure. By calculating the average RMSD for each receptor structure, the most favorable structure is identified based on the lowest average RMSD value. Subsequently, to rectify any potential structural incompleteness, the selected receptor structure is subjected to hydrogen addition using OpenBabel [20]. This step aims to address any deficiencies or missing hydrogen atoms, thereby enhancing the overall completeness and accuracy of the receptor structure. The meticulous selection and preparation of the receptor structure in Vina-GPU 2.1 significantly contribute to the subsequent accuracy and reliability of the virtual screening process. By employing the cross-docking method and addressing structural incompleteness, Vina-GPU 2.1 aims to optimize the receptor structure and improve the efficacy of structure-based virtual screening.

#### 2) Pocket Prediction

The second step is to predict the binding pocket of the optimized receptor structure. Instead of directly obtaining binding pocket from the crystal structure, we utilize COACH-D[14] to identify the binding pocket with an improved accuracy. The COACH-D method uses TM-SITE that structurally compares the uploaded receptor to the templates in the database with a balanced approach on the (global) entire structure and the (local) binding site region. Specifically, we upload the optimized receptor structure (from step 1) to the COACH-D server (https://yanglab.nankai.edu.cn/COACH-D/), which gives out five predictions of the pocket and the indexes of residues accordingly. We use the top 1 prediction ranked by C-score[14] and calculate the pocket size and center by Equation.4 and Equation.5,

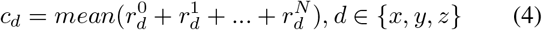

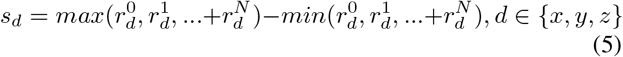

where *c* and *s* represent the center and the size of the pocket, *r*^*i*^ means the *i*-th residue of the given protein and {*x, y, z*} stands for the three orthogonal dimensions in the Cartesian coordinates. The operations *mean*(), *max*() and *min*() calculate the mean, the maximum and the minimum of the inputs, respectively.

#### 3) Ligand Optimization

The third step is to optimize the structure of the small molecules obtained from large data bases because of the ill-consideration when using classic method like OpenBabel [20] (adopted in Vina-GPU 2.0). Gypsum-DL [12] is a free and robust program that takes ionization, tautomeric, and ring-conformational variants into consideration and it offers prominent accuracy improvement in field of virtual screening. In this step of Vina-GPU 2.1, we use Gypsum-DL to generate the 3D structures of the molecules from SMILES format to SDF format and then perform another translation from SDF format to PDBQT format with the rdkit2pdbqt.py script.

## III. RESULTS AND DISCUSSION

### A. Experiment Settings

#### Protein Targets

We choose three important and widely studied proteins to conduct our experiments in this study. The first one is the RIPK1 protein, which serves as a signal transducer in various cellular processes, including cell proliferation, differentiation, and cell death. The second one is the RIPK3 protein, which is primarily known for its role in necroptosis, a form of programmed cell death that occurs when apoptosis is inhibited [21]. Both of RIPK1 and RIPK3 play the acritical roles in regulating cell death and inflammation, which are essential for maintaining tissue homeostasis and immune responses. Dysregulation of the two receptors has been implicated in the pathogenesis of various diseases, including cancer, neurodegenerative disorders, and inflammatory diseases [22]. The third one is the AmpC *β*-lactamase, which is an enzyme produced by certain bacteria that confers resistance to a wide range of *β*-lactam antibiotics, including penicillins, cephalosporins, and carbapenems. The emergence and spread of AmpC *β*-lactamase-producing bacteria has become a major public health concern, as it limits the effectiveness of many antibiotics and complicates the treatment of infections caused by these bacteria.

#### Compound Library

We choose to use two widely used molecular databases to conduct virtual screening in this study. The first one is the DrugBank [23] library, which is a comprehensive database that provides information on drugs, their targets, and their mechanisms of action. We select 9125 molecules with its structures from DrugBank at https://go.drugbank.com/releases/latest#. The second one is the Selleck [24] Library, which includes a wide range of small molecule compounds, including inhibitors, agonists, antagonists, and activators of various cellular targets. We obtained 5148 molecules from the L460-TargetMol-Natural Compound Library and the L1400-Selleck-Natural-Product-Library-131cpds plate. Our dataset is available at https://github.com/DeltaGroupNJUPT/Vina-GPU-2.1/tree/main/experimentdataset.

#### Evaluation Metrics

We use the enrichment factor (*EF*) and the number of hits compound (*Hit*)as the evaluation metrics in our virtual screening experiments. The definition of *EF*_*x*%_ and *Hit*_*x*%_ are given in Equation 6 and Equation 7, which include the number of active compounds (*Hit*_*s*_) within the top *x*% compounds (*N*_*s*_) and the number of total active compounds (*Hits*_*t*_) within the total number of database (*N*_*t*_).

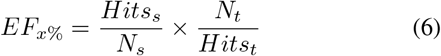

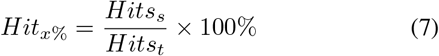

#### Active compound

We manually collected active compounds for RIPK1, RIPK3 and AmpC *β*-lactamase from ChEMBL [25] website with the 1 *uM* activity threshold. All the active compounds are provided in https://github.com/DeltaGroupNJUPT/Vina-GPU-2.1/tree/main/experimentdataset.

#### Experimental settings

The experimental settings for a total of 7 methods are given in Table I, in which the parameters are chosen carefully according to the Vina-GPU [6] and Vina-GPU 2.0 [5]. The search depth in AutoDock Vina-GPU 2.1 is set heuristically by Equation 8, which is 1.5 times larger than those in Vina-GPU [6] and Vina-GPU+[5].

**TABLE 1.**
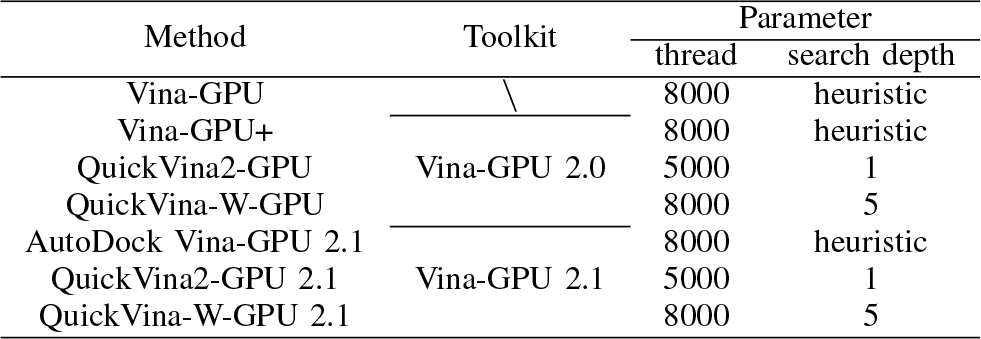
Experimental settings.

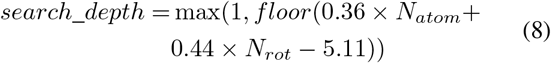

For a fair comparison, we utilize the same experiment parameters (Table I) and computation resources under the same compound library. For the DrugBank database (in III C), we used Intel (R) Core (TM) i9-10900K CPU @ 3.7 GHz with the exhaustive use of 20 CPU cores as the CPU device, and Nvidia Geforce RTX 3090 (OpenCL v3.0) under single-precision floating-point format (FP32) as the GPU device. For the Selleck database (in III D), we used Intel (R) Core (TM) i7-12700H CPU @ 2.7 GHz with the exhaustive use of 20 CPU cores as the CPU device and NVIDIA Geforce RTX 3060 Laptop (OpenCL v3.0) under single-precision floating-point format (FP32) as the GPU device.

### B. Better Docking on benchmark dataset

The docking benchmark is conducted on the AutoDock-GPU 140 dataset, which includes 140 complexes that are well pre-processed according to the original literatures [26], [27], [28]. Therefore, no structure optimizations are needed for these complexes since we only benchmark the docking performance (instead of virtual screening) of our methods.

To validate the time consumptions of each runtime phase of AutoDock Vina, we randomly selected six complexes from the AutoDock-GPU 140 dataset and used AutoDock Vina to conduct the molecular docking. The time consumptions of each phase were given in Table II. It is obvious that the BFGS optimization phase is the most time-consuming one that takes over 82% of the total runtime.

**TABLE 2.** Time (in seconds) consumption of different phases of AutoDock vINA ALGORITHM ON DIFFERENT COMPLEXES.

| PDB id | Input files calculation | Grid cache computing | BFGS optimization | Fine-grained optimization |
| --- | --- | --- | --- | --- |
| 5tim | 0.014(0.9%) | 0.232(16.1%) | 1.193(82.7%) | 0.003(0.2%) |
| 1ac8 | 0.0133(0.7%) | 0.330(16.3%) | 1.674(82.7%) | 0.008(0.40%) |
| 2bsm | 0.0141(<0.1%) | 0.503(1.20%) | 40.751(98.6%) | 0.045(0.1%) |
| 2bm2 | 0.0131(<0.1%) | 0.440(<0.1%) | 48.569(>99.0%) | 0.060(0.1%) |
| 4er4 | 0.0142(<0.1%) | 0.434(<0.1%) | 2356.765(>99.9%) | 0.396(<0.1%) |
| 1jyq | 0.0140(<0.1%) | 0.383(<0.1%) | 472.151(>99.9%) | 0.177(<0.1%) |

Then, we conducted a docking benchmark for the Vina-GPU 2.0 and the proposed Vina-GPU 2.1 methods on the AutoDock-GPU 140 dataset [29] and the results are given in Table III. Top-1 and Top-5 means the top 1 and the top 5 of the docking results, respectively. There is no obvious difference between the methods in Vina-GPU 2.1 and the corresponding method in Vina-GPU 2.0 in terms of the RMSD (between output ligand and the x-ray ligand) and the docking score. Vina-GPU+ and AutoDock Vina-GPU 2.1 show better docking accuracy among these methods, while QuickVina2-GPU 2.1 and QuickVina-W-GPU 2.1 show better docking speed. The averaged runtime of each sample is largely reduced by 38.3%, 85.5% and 71.9% for AutoDock Vina-GPU 2.1, QuickVina2-GPU 2.1 and QuickVina-W-GPU 2.1. Thus, the three methods of Vina-GPU 2.1 achieve 1.62*×*, 6.89*×* and 3.56*×* acceleration.

**TABLE 3.** Docking benchmark on the aUTOdOCK-GPU 140 DATASET.

| Method | Top-1 RMSD(Å) |  | Top-5 RMSD(Å) |  | Average Top-1 Score (kcal/mol) | Average Top-5 Score (kcal/mol) | Average Runtime (s) |
| --- | --- | --- | --- | --- | --- | --- | --- |
|  | % <2 | average | % <2 | average |  |  |  |
| Vina-GPU+ | 0.56±0.01 | 3.11±3.26 | 0.71±0.00 | 3.86±3.21 | -7.59±1.63 | -7.55±1.62 | 5.46 |
| AutoDock Vina-GPU 2.1 | 0.57±0.01 | 3.15±3.41 | 0.70±0.01 | 3.88±3.23 | -7.62±1.62 | -7.59±1.60 | 3.37 |
| QuickVina2-GPU | 0.46±0.01 | 3.84±3.54 | 0.61±0.01 | 5.45±3.23 | -6.75±1.72 | -6.58±1.54 | 2.00 |
| QuickVina2-GPU 2.1 | 0.45±0.01 | 3.81±3.52 | 0.58±0.02 | 4.75±3.42 | -6.96±1.64 | -6.88±1.57 | 0.29 |
| QuickVina-W-GPU | 0.54±0.01 | 3.20±3.34 | 0.69±0.02 | 4.98±3.19 | -7.36±1.70 | -7.18±1.56 | 3.67 |
| QuickVina-W-GPU 2.1 | 0.54±0.02 | 3.37±3.53 | 0.66±0.02 | 4.20±3.35 | -7.35±1.65 | -7.30±1.61 | 1.03 |

### C. Better Virtual Screening from DrugBank Library

The virtual screening runtime results of three receptors on the DrugBank library is shown in Figure 2, where AutoDock Vina-GPU 2.1, QuickVina2-GPU 2.1 and QuickVina-W-GPU 2.1 achieve the highest screening speed and obtain 2.91, 11.15 and 4.23 accelerations (averaged across three targets on the DrugBank database) when comparing to Vina-GPU+, QuickVina2-GPU and QuickVina-W-GPU, respectively. The QuickVina2-GPU 2.1 method achieves the fastest docking speed on the DrugBank library due to the I-check mechanism that reduces redundant iterations. Although QuickVina-W-GPU 2.1 adopts both I-check and G-check, its computational cost is relatively higher than that of QuickVina2-GPU 2.1. The local and global buffers of G-check bring additional overhead that makes QuickVina-W-GPU 2.1 inefficient. The virtual screening accuracy is illustrated in Figure 3. In general, our AutoDock Vina-GPU 2.1, QuickVina2-GPU 2.1 and QuickVina-W-GPU 2.1 methods achieve superiors screening accuracy when compared to Vina-GPU+, QuickVina2-GPU and QuickVina-W-GPU, respectively. In average, Vina-GPU 2.1 obtains 321%/312%/209% and 319%/313%/210% improvement in Hit (1% / 5% / 10%) and EF (1% / 5% / 10%), respectively, when compared to Vina-GPU 2.0. In addition, our AutoDock Vina-GPU 2.1 method achieves the highest average performance on the Hit and EF metrices. Notably, although QuickVina2-GPU 2.1 achieves the highest docking speed, the virtual screening accuracy is degraded. For the DrugBank library, we recommend using QuickVina2-GPU 2.1 for users that require a fast docking speed and AutoDock Vina-GPU 2.1 for users that prefer a high screening accuracy.

**Fig. 2.**
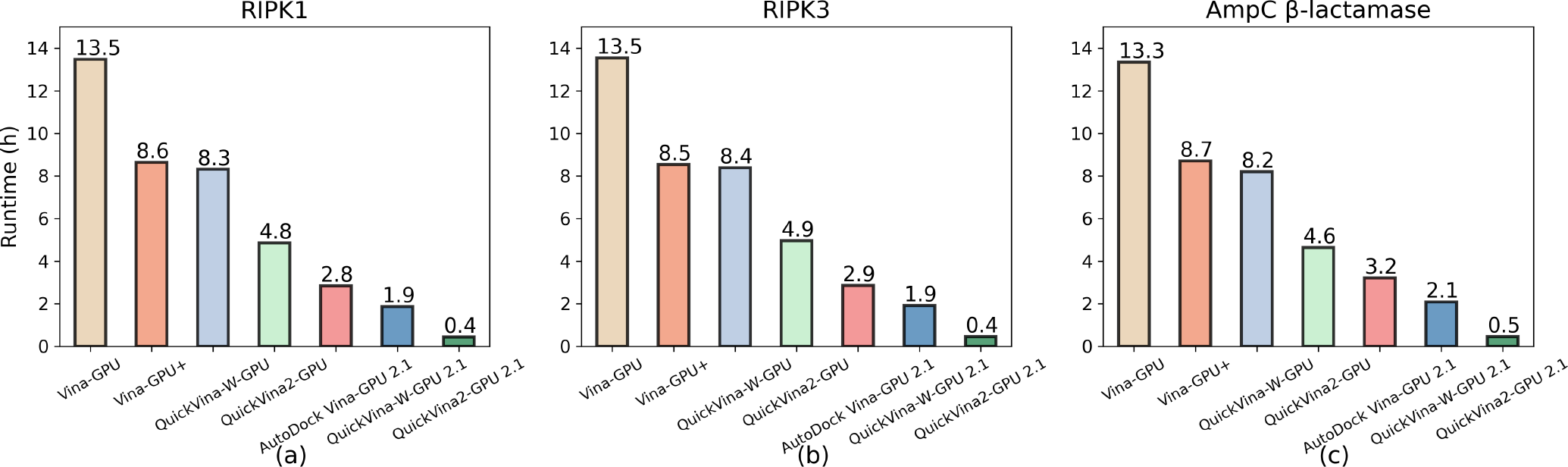
Faster virtual screening from DrugBank library by seven docking methods on three targets (RIPK1, RIPK3 and AmpC *β*-lactamase).

**Fig. 3.**
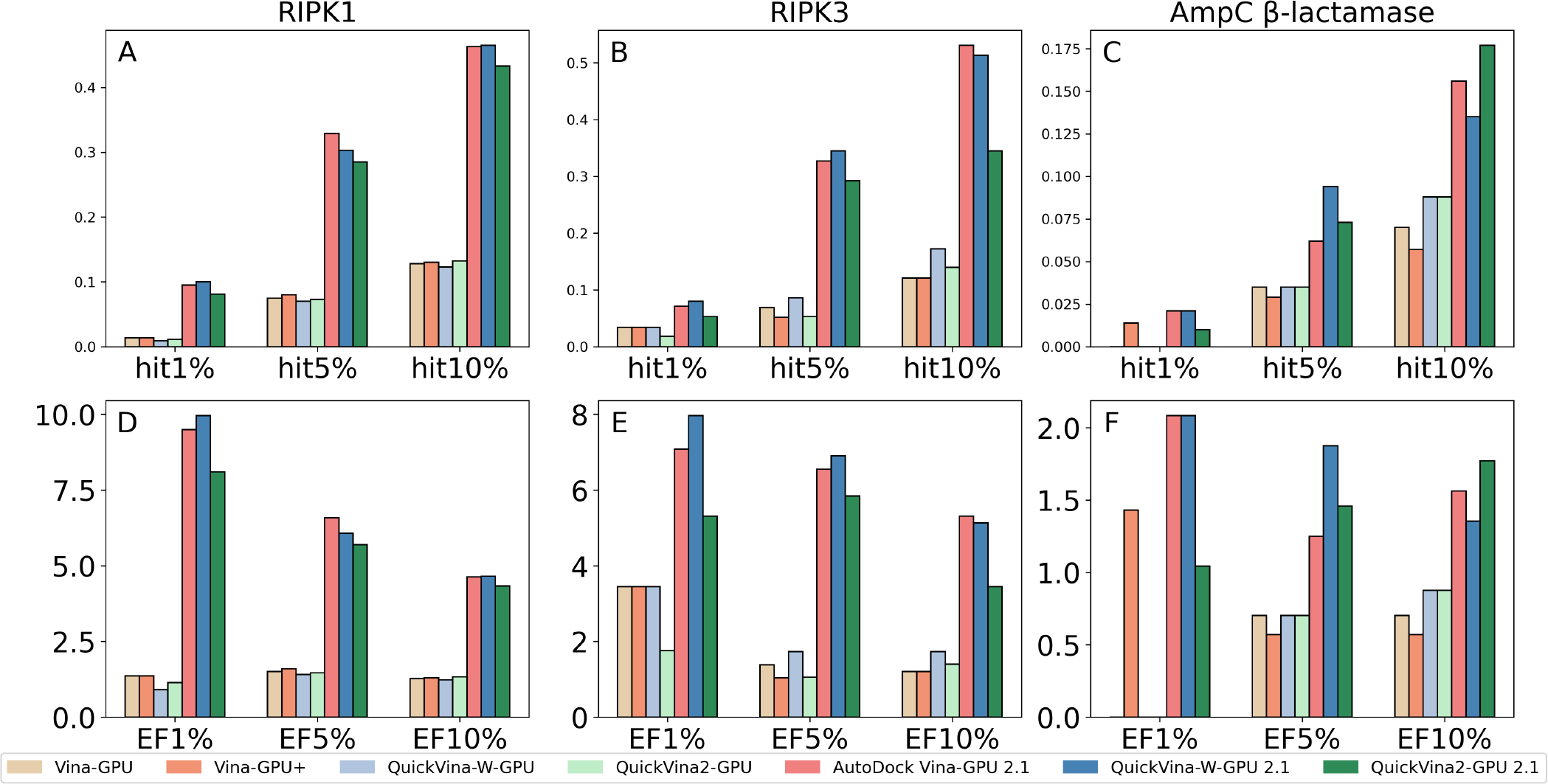
Comparison of screening accuracy by seven docking methods from DrugBank library on three protein targets.

### D. Better Virtual Screening from Selleck Library

The virtual screening runtime results of three receptors on the Selleck library is shown in Figure 4, where AutoDock Vina-GPU 2.1, QuickVina2-GPU 2.1 and QuickVina-W-GPU 2.1 achieve the highest screening speed and obtain 1.44, 5.53 and 4.60 accelerations (averaged across three targets on the Selleck database) when comparing to Vina-GPU+, QuickVina2-GPU and QuickVina-W-GPU, respectively. The QuickVina2-GPU 2.1 obtains the highest docking speed across all seven methods due to the I-check mechanism. The virtual screening accuracy is illustrated in Figure 5. In general, our AutoDock Vina-GPU 2.1, QuickVina2-GPU 2.1 and QuickVina-W-GPU 2.1 methods achieve superiors screening accuracy when compared to Vina-GPU+, QuickVina2-GPU and QuickVina-W-GPU, respectively. In average, Vina-GPU 2.1 obtains 328%/440%/231% and 365%/445%/231% improvement in Hit (1% / 5% / 10%) and EF (1% / 5% / 10%), respectively, when compared to Vina-GPU 2.0. The accuracy improvement indicates that the structure optimization (II D) contributes well to the virtual screening of the Selleck library. In addition, our AutoDock Vina-GPU 2.1 method achieves the highest average performance on the EF and the Hit metrics. For the Selleck library, we also recommend using QuickVina2-GPU 2.1 for users that require a fast docking speed and AutoDock Vina-GPU 2.1 for users that prefer a high screening accuracy.

**Fig. 4.**
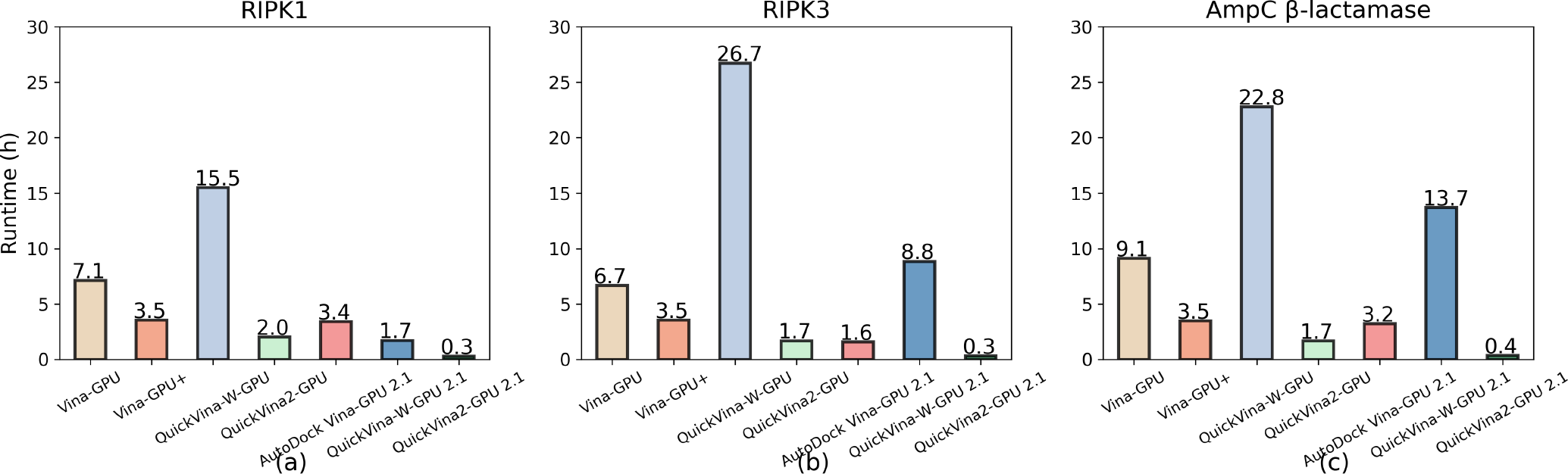
Faster virtual screening from Selleck libraryby seven docking methods on three targets (RIPK1, RIPK3 and AmpC -lactamase).

**Fig. 5.**
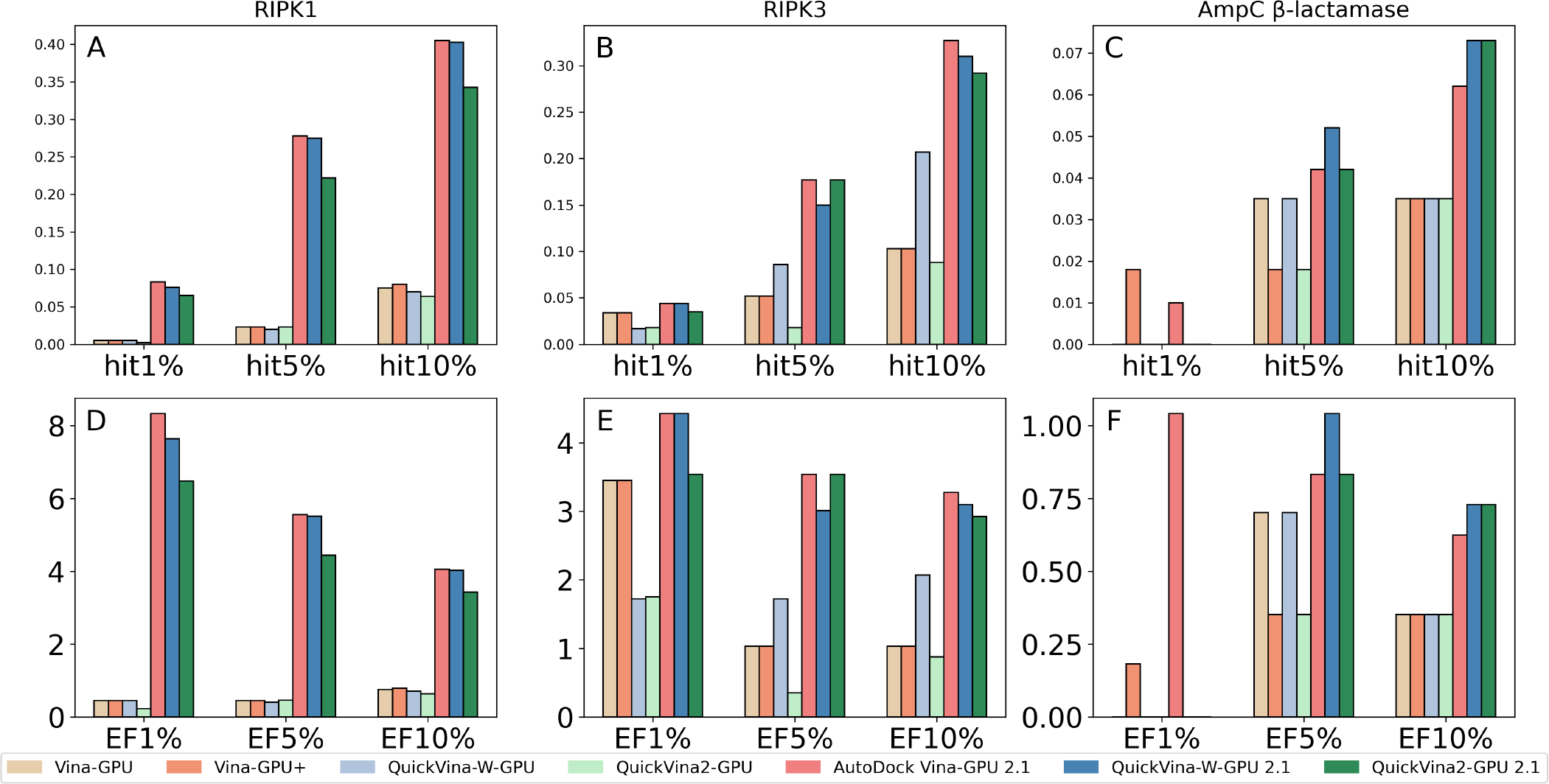
Comparison of screening accuracy by seven docking methods from Selleck library on three protein targets.

### E. Ablation study on virtual screening

#### Screening accuracy

We conducted an ablation study to illustrate the benefit of each structure optimization on the screening accuracy in Figure 6. The experiment is targeted on the RIPK1 receptor with the DrugBank dataset, and all active molecules are gathered from ChEMBL website. In Figure 6, “cross-docking”, “COACH-D” and “pyg” represent the three structure optimizations (as introduced in II D) in the Vina-GPU 2.1 method. “lowest res” represents using the experimental determined protein structures with the lowest resolution. “plugin” represents using the Pymol [30] plugin to determine the docking pocket with the experimental binding ligand. “babel” represents using OpenBabel to obtain ligand structures from the database. In general, each of the three structure optimizations contribute to the improvement of the Hit (1% / 5% / 10%) and EF (1% / 5% / 10%). Few exceptions occur at the COACH-D optimization due to the randomness of the Vina-GPU 2.1 method and the quality of the active compounds collected from the ChEMBL website.

**Fig. 6.**
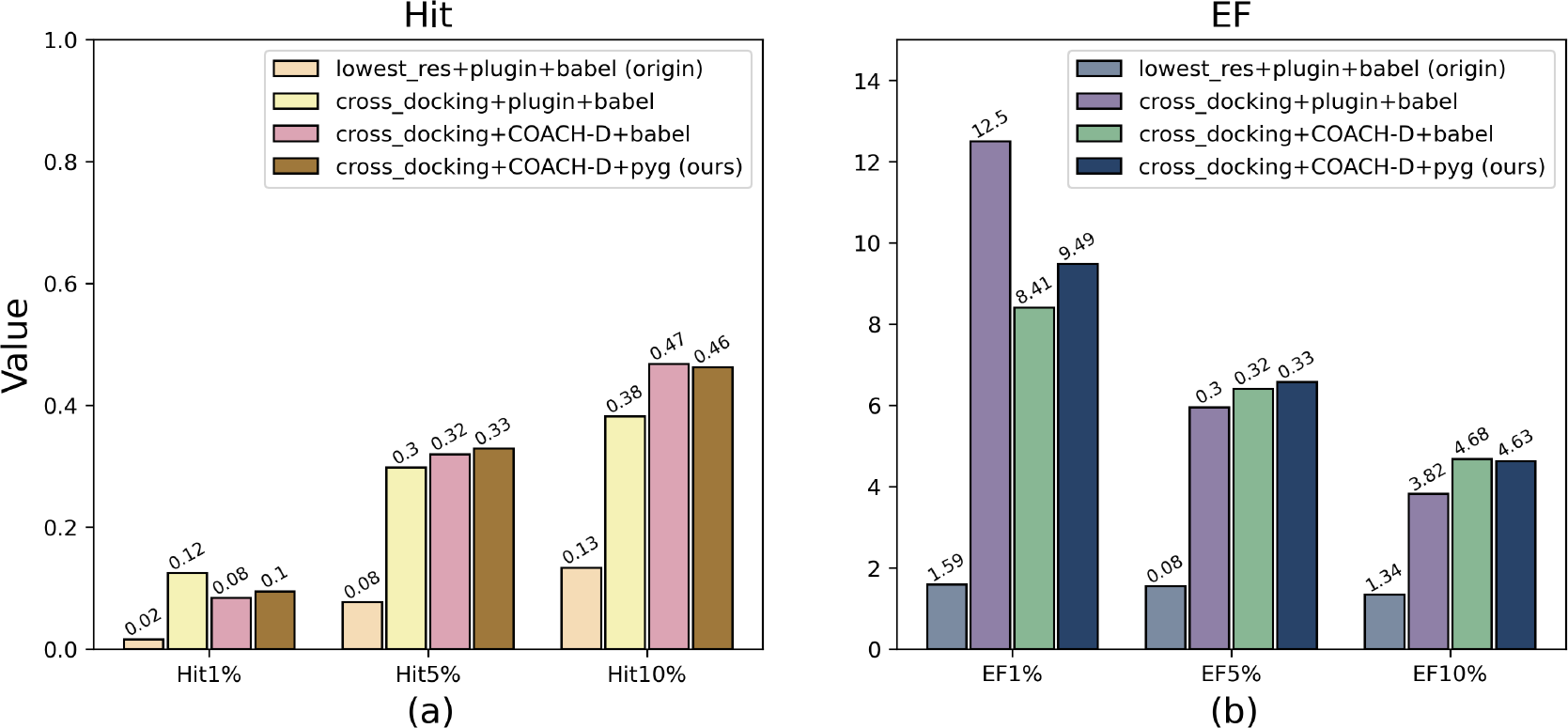
Ablation study on screening accuracy. “lowest res” selects receptor structure with the lowest crystal resolution. “plugin” uses Pymol plugin for automatic binding box determination. “babel” employs OpenBabel for ligand conformation conversion from 2D to 3D. “cross docking” utilizes cross docking method for receptor structure selection. “COACH-D” relies on the COACH-D server for binding pocket determination. “pyg” uses Gypsum-DL for ligand conformation conversion.

#### Screening runtime

We conducted an ablation study to illustrate the benefit of each sampling optimization on the screening runtime in Figure 7. The experiment is targeted on the RIPK1 receptor with the DrugBank dataset. In Figure 7, “origin” represents the AutoDock Vina-GPU 2.1, QuickVina2-GPU 2.1 and QuickVina-W-GPU 2.1 that proposed in our Vina-GPU 2.1 method. “RILC-BFGS” and “GCS” represent the two efficient sampling methods proposed in II B and II C). As shown in Figure 7, each of the two sampling methods contributed to the acceleration of the screening runtime. In average, the proposed RILC-BFGS and GCS method achieved 47.27% and 53.41% reduction on the virtual screening runtime.

**Fig. 7.**
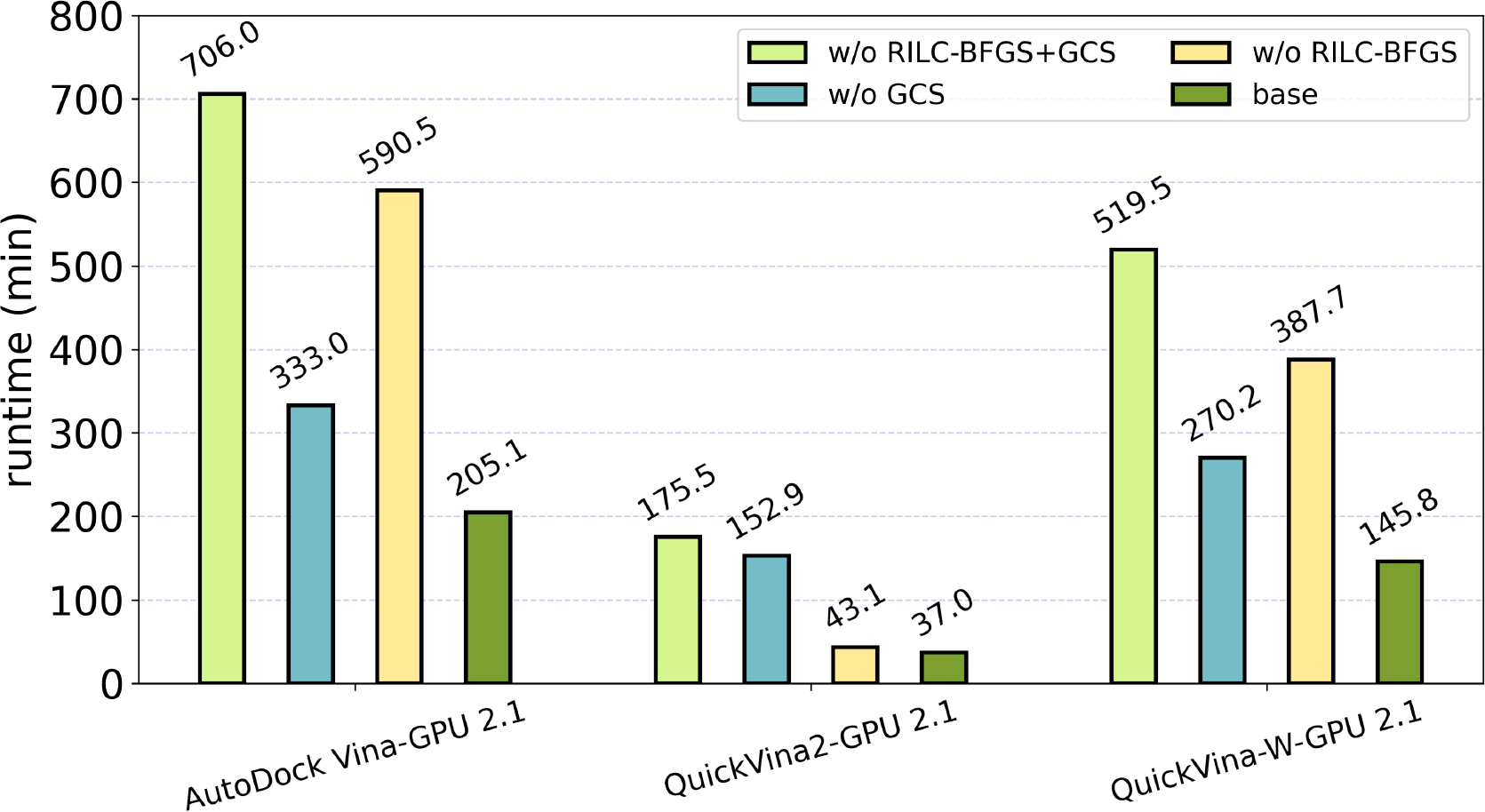
Ablation study on screening speed. “RILC-BFGS” and “GCS” refers to the two methods proposed in this study. “base” refers to the Vina-GPU 2.1.

### F. Guideline for Usage

In this work, We introduce our Vina-GPU 2.1 as a robust and highly effective toolkit that demonstrates its unwavering reliability, making it an exemplary choice for researchers and practitioners in the field. To improve user experience of our Vina-GPU 2.1 toolkit, we open-sourced Vina-GPU 2.1 toolkit at Github with detailed usage guideline that helps experts and non-experts to easily get started to conduct virtual screening. The guideline helps users to build Vina-GPU 2.1 from source (including the OpenCL/CUDA environment and the Boost library) step by step and provides detailed instructions to run Vina-GPU 2.1 with the typical (docking) mode and the virtual screening mode. The guideline also entails the details of the structure optimizations, providing exact link/reference that embedded in the Vina-GPU 2.1. Since the development of Vina-GPU, we have consistently received kind feedbacks from users to improve user experience and reduce runtime errors.

Until now, we have fixed dozens of bugs and embedded several important functionalities such as log outputs, error detection, runtime profiling etc. into our toolkit. For many developers that are more familiar with the CUDA programming language, we also provide an open-sourced CUDA version of Vina-GPU at https://github.com/Glinttsd/Vina-GPU-CUDA.

## IV. CONCLUSION

Speed and accuracy are crucial factors in modern drug discovery, and we have made significant advancements in these areas with our Vina-GPU and Vina-GPU 2.0. Building upon this success, we introduce Vina-GPU 2.1, which pushes the boundaries even further. Vina-GPU 2.1 incorporates the innovative RILC-BFGS algorithm, which integrates several efficient algorithms to reduce computational complexity. This optimization enhances the screening speed while maintaining high accuracy. By leveraging these structure optimization methods, Vina-GPU 2.1 outperforms both Vina-GPU and Vina-GPU 2.0 in terms of speed and accuracy. With its exceptional screening speed and accuracy, Vina-GPU 2.1 becomes a powerful toolkit in the modern drug discovery pipeline. Researchers can rely on this tool to expedite the screening process and identify potential drug candidates more efficiently. In future endeavors, we will concentrate on optimizing the RILC-BFGS algorithm specifically for GPU architectures and leveraging the capabilities of machine learning algorithms to enhance the quality of docked conformations.

## V. acknowledgement

This work was supported by the National Natural Science Foundation of China (62371245, 61872198). Thanks to Prof. Jieping Ye and Mr. Junlong Liu of Alibaba Damo Academy for their guidance and assistance. Thanks to Alibaba Damo Academy for their cooperation and support of this project. Thanks to Hao Zhang and Chenyu Wang for their support of this project.

